# (+)-*trans*-Cannabidiol is a CB_2_ receptor agonist

**DOI:** 10.64898/2026.05.25.727077

**Authors:** Pedro Bans Burtchaell, Marina Junqueira Santiago, Chenxi Wang, Mehdi Hagdoost, Evie J. M. Clay, Dani Mohnot, Mark Connor

**Author notes:** Author for correspondence: Mark Connor, Macquarie Medical School, Level 1, 75 Talavera Rd, Macquarie University, NSW, 2109., MC.

## Abstract

(*−*)*-trans*-Cannabidiol ((−)-CBD) is a principal phytocannabinoid from *Cannabis sativa*. (−)-CBD has complex pharmacology but is a relatively weak inhibitor of CB1 and CB2 receptor signalling. Cannabidiol has two chiral centres and thus four stereoisomers. (+)-*trans*-CBD ((+)-CBD) has a higher affinity than (−)-CBD at CB1 and CB2, but its pharmacodynamic effects at these receptors are incompletely described. We examined the activity of (+)-CBD at human CB1 and CB2 receptors using a fluorescence-based assay of membrane potential in AtT20 cells stably expressing CB1 or CB2 receptors. (+)-CBD produced a rapid, concentration-dependent hyperpolarization in CB2-expressing cells (*p*EC50 6.63 ± 0.08) with a maximal effect ∼90% of the response to CP55940. The CB2 response was blocked by pertussis toxin pretreatment and competitively inhibited by the CB2 antagonist AM630 (Schild slope 1.1 ± 0.1). (+)-CBD was a low-efficacy, low-potency CB1 agonist and inhibited somatostatin-receptor effects at high concentrations (10-30 µM). It had no effect on the membrane potential of AtT20 wild-type cells. In *silico* modelling of ligand interactions with CB2 indicated that (+)-CBD but not (−)-CBD formed an H-bond with Ser285, a residue crucial for agonist activation of CB2. Our data suggests (+)-CBD acted as a CB2 agonist via the orthosteric binding site on the receptor. Synthetic CBD, including (+)-CBD, has previously been administered in clinical trials, presumably without consideration of its potential CB2 agonist activity. Given the relative safety of (−)-CBD in people, (+)-CBD may be a useful drug to explore CB2-sensitive disease states, should it prove similarly safe.

## Introduction

Cannabidiol is one of the major biologically active constituents of *Cannabis*, and is a drug used in the treatment of some seizure disorders, multiple sclerosis and chronic pain (1). The mechanisms of action of CBD in these conditions remain poorly defined, although among the dozens of potential targets for CBD identified *in vitro*, interactions with ion channels (2,3,4), and G protein-coupled receptors (5,6,7) provide potentially plausible sites of action. CBD also affects drug-metabolising enzymes (8,9), leading to pharmacokinetic drug-drug interactions which may contribute to its clinical effects. CBD is generally described as a negative allosteric modulator of CB_1_ receptors (10, 11) while there are contrasting reports of its activity at CB_2_ (11,12,13,14).

CBD has 2 chiral centres, and thus 4 potential stereoisomers (15). (−)-*trans*-CBD is to date the only isomer found in *Cannabis*, which is in contrast to other chiral cannabinoids, as all 4 isomers of Δ^9^-tetrahydrocannabinol (THC) have been identified in some strains of *Cannabis* (16,17), as have both (−) and (+)-cannabichromene (CBC) (16,18). (+)-CBD can be readily accessed through synthetic routes (15,19,20), and a small number of studies have reported biological activities of (+)-CBD distinct from those of (−)-CBD. These include antagonism of CB_1_-mediated responses at sub-µM (+)-CBD concentrations in mouse neurons *in vitro*, activation of human S1P_1_ and S1P_3_ receptors in HEK293 cells (21) and a substantially higher affinity of (+)-CBD for CB_2_ receptors than CB_1_ receptors (19).

CBD is currently being trialled in people to treat many conditions (22), presumably in part because of its relatively good safety profile and the strong anecdotal evidence about the efficacy of *Cannabis* for an extraordinarily wide range of ailments. It is not always clear what molecular form of CBD is used in the trials – plant derived CBD is likely to be (−)-trans CBD, but if synthetic CBD is being used, then there is a possibility that (+)-CBD is being administered to patients as either a pure substance or part of a racemic mixture. Use of pure (+)-CBD was reported in one trial (23); however, in other trials using synthetic CBD, the drug’s chirality is not reported. Given the importance of chirality in cannabinoid action (17,24,25), and the moderate affinity of (+)-CBD for CB_2_ receptors (19), we explored the activity of (+)-trans CBD at cannabinoid receptors *in vitro*, finding that it has robust agonist activity at CB_2_ receptors.

## Methods

### Cell Culture

AtT20-FlpIn cells stably expressing 3x haemagglutinin-tagged human CB_1_ or CB_2_ (AtT20-CB_1_; AtT20-CB_2_) (26) or untransfected AtT20-FlpIn cells (AtT20-WT; first described in (27)) were cultured in DMEM medium containing 10% FBS and 100 units penicillin-streptomycin (P/S) in a humidified incubator at 37 °C with 5% CO_2_. Media for AtT20-CB_1_, and AtT20-CB_2_ cells also contained the selection antibiotic hygromycin (80 µg.mL^−1^). Cells were fed every 3-4 days and passaged when they had reached approximately 80% confluence. Cells were routinely tested for mycoplasma using MycoAlert detection kit (Lonza) and were always negative.

For experiments, Leibovitz’s L-15 medium supplemented with 1% FBS, 100 units P/S, and 15 mM glucose was used. Cells were cultured overnight in a humidified incubator at 37 °C, ambient air (no added CO_2_).

### Membrane Potential Assay

Changes in membrane potential were measured using a proprietary membrane potential assay kit (MPA Blue #R8034, Molecular Devices, San Jose, CA, USA) as described previously (28). AtT20-CB_1_, AtT20-CB_2_, or AtT20-WT cells were incubated overnight in 90 µL of L-15-supplemented medium in 96-well black-walled, clear-bottom plates (Corning). The same volume (90 µL) of MPA dye dissolved in a modified Hank’s Buffered Saline Solution (HBSS) was added to the cells at 50% of the manufacturer’s recommended concentration, then placed in a FlexStation 3 (Molecular Devices) for at least 60 minutes at 37 °C. For single additions, drugs were dissolved in HBSS supplemented with 0.1% bovine serum albumin (Sigma-Aldrich) and were added in a volume of 20 µL (for a final volume of 200 µL) after a baseline recording of 60-120s; for double additions the second addition was 22 µL. Fluorescence was recorded every 2 seconds with excitation at 530 nm and emission at 565 nm (550 nm cut-off).

Drug effects are reported as the peak change in fluorescence following drug addition, expressed as a percentage of the pre-drug baseline. Each column included one well in which only vehicle-control (buffer + DMSO) was added; the changes in fluorescence produced by these vehicle-control additions were subtracted from the experimental traces. The final DMSO concentrations in the well were between 0.1% (single addition) and 0.2 % (double addition). Concentration-response parameters were derived from at least six independent experiments, performed in duplicate and then fitted to a 4-parameter logistic equation in PRISM (version 10.3.1, GraphPad Software, Boston, MA, USA). Values for pA2 and Schild slope were calculated for individual experiments in PRISM, and the values were averaged. Data is presented as mean ± SEM of at least six independent replicates; exact values are noted in the text. Statistical comparisons of untransformed values were made using an unpaired Student’s t-test or an ordinary one-way ANOVA followed by Dunnett’s multiple-comparisons test versus control (PRISM).

### In silico modelling

Structural templates of the CB_2_ receptor were retrieved from the Protein Data Bank. A total of seven X-ray and cryo-EM structures (PDB IDs: 6PT0 (29), 8GUQ, 8GUR, 8GUT 6KPF, 6KPC (30) and 5ZTY (31)) were evaluated. Among these, the 8GUR structure was selected as it provided the most accurate activity predictions for a benchmark set of active and decoy CB_2_ ligands (32). Protein preparation for molecular docking was performed using UCSF Chimera’s (RBVI, University of California, San Francisco, CA) Dock Prep protocol. This included the removal of solvents, unbound ions, and the co-crystallized ligand. Incomplete side chains were reconstructed using the Dunbrack 2010 rotamer library (33), while missing hydrogen atoms were added and partial charges assigned using the AMBER ff99bsc0 force field (34). Energy minimization of the receptor structure was omitted, as it did not enhance model performance on the active–decoy dataset. The binding site and docking grid were defined using AutoDockTools4 (35) with the following grid centre coordinates: X = 135, Y = 145, Z = 168. The 3D structures of (−)-CBD, (+)-CBD, and CP55940 were downloaded from PubChem (36). and geometry-optimized using the MMFF94s force field (37).

Molecular docking was performed using AutoDock Vina 1.2 (38). During the docking method assessment, three scoring functions (Vina, Vinardo (39), and a recently reported custom empirical set (40)) were evaluated. The Vinardo scoring function, which demonstrated superior performance in distinguishing actives from decoys, was selected for subsequent docking analyses. A grid box size of 25 × 25 × 25 Å³ and an exhaustiveness value of 100 were applied to all docking calculations, as increasing exhaustiveness did not yield significant improvements in docking scores. For each ligand, the top three best-scoring poses were retained for further analysis and comparison.

### Drugs and HBSS solution

(−)-CBD was from the National Measurement Institute (North Ryde, Australia). (+)-CBD (2-[(1S,6S)-3-methyl-6-(1-methylethenyl)-2-cyclohexen-1-yl]-5-pentyl-1,3-benzenediol), AM630, SR141716A and CP55940 were from Cayman Chemical. ML297 was from Sigma-Aldrich. Somatostatin was from AUSPEP (Melbourne, Australia). Pertussis Toxin was from HelloBio (Bristol, UK).

To confirm that the (−)-CBD and (+)-CBD we used were indeed optical isomers, we obtained their respective spectra using circular dichroism (CD, Supplementary Figure 1). Briefly, the circular dichroism and absorbance spectra of the samples were determined using the Jasco J-1500 spectropolarimeter (Hachioji, Japan). Conditions were optimised prior to final readings. Samples of (+)-CBD and (−)-CBD were diluted to 0.1mg.mL^−1^ in 100% ethanol and a pathlength of 0.1mm was used. For each sample, 32 runs were carried out, and each curve represents the average of the 32 runs. The pure ethanol control reading was subtracted from the sample readings, and these blank-corrected readings used to produce the spectra. Measurements were carried out at room temperature. Measurements were made twice with different samples of (−)-CBD and (+)-CBD, with similar results.

HBSS was prepared by mixing (mM) NaCl 145, HEPES 22, Na_2_HPO_4_ 0.338, NaHCO_3_ 4.17, KH_2_PO_4_ 0.441, MgSO_4_ 0.407, MgCl_2_ 0.493, glucose 5.56, and CaCl_2_ 1.26. The solution pH was adjusted to 7.4, and osmolarity was 315 ± 15 mOsmol.

## Results

### The effects of (+)-CBD at AtT20-CB_2_, CB_1_ and WT

Application of (+)-CBD produced a rapid, concentration-dependent hyperpolarisation of AtT20-CB_2_ cells with a *p*EC_50_ of 6.63 ± 0.08 (n_H_ 1.3 ± 0.3), and maximum change in fluorescence of 24.7 ± 0.9 % (Figure 1A and 1D, black closed-circle; n = 7). In comparison, the non-selective cannabinoid receptor agonist CP55940 hyperpolarised AtT20-CB2 cells with a *p*EC_50_ of 7.48 ± 0.08 (n_H_ 1.2 ± 0.2) to a maximum of 27.2 ± 1.2 % (Figure 1A and 1D, open-diamonds; n = 6).

**Figure 1.**
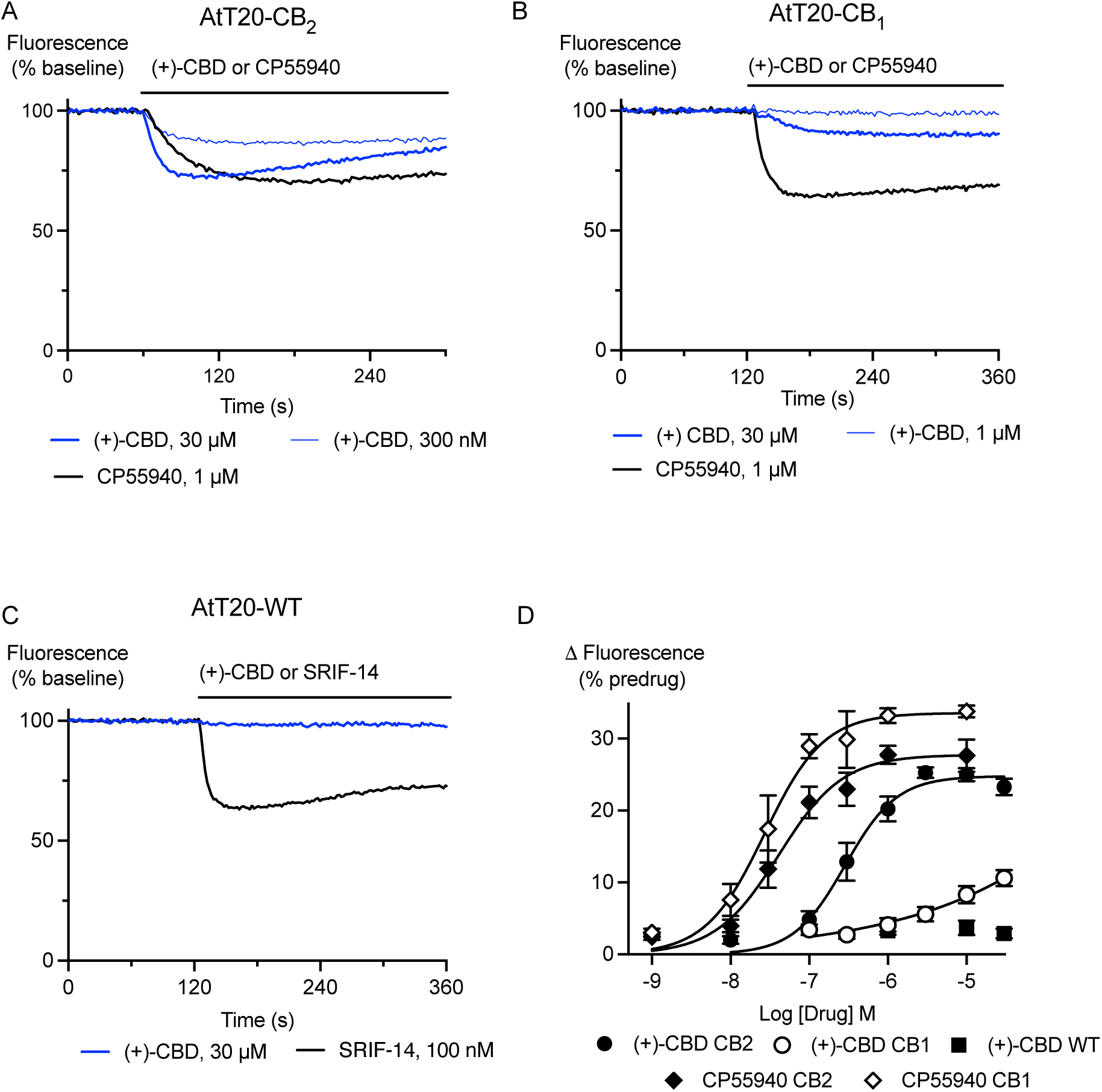
(+)-CBD hyperpolarises AtT20-CB_2_ and AtT20-CB_1_ cells. AtT20 cells expressing CB_2_, CB_1_ or no cannabinoid receptors (WT) were challenged with (+)-CBD and CP55940. A fluorescent membrane potential dye was used to measure the response. **A)** Representative traces showing that application of (+)-CBD (300 nM, 30 µM) and CP55940 (1 µM) to AtT20-CB_2_ cells produced a sustained decrease in fluorescence, consistent with hyperpolarisation of the cells **B)** Representative traces showing that application of (+)-CBD (1 µM, 30 µM) to AtT20-CB_1_ cells produced much smaller change in fluorescence than CP55940 (1 µM). **C)** Representative traces showing that (+)-CBD (30 µM) did not produce a change in fluorescence in AtT20-WT cells, but somatostatin-14 (SRIF-14, 100 nM) did. **D)** Concentration-response curves for (+)-CBD, and CP55940 showing the maximum change in fluorescence produced by each drug in AtT20-CB_1_, -CB_2_ and -WT cells. Each point represents the mean ± SEM of at least 6 independent experiments, performed in duplicate.

High concentrations of (+)-CBD produced a modest hyperpolarisation of AtT20-CB_1_ cells, with only 10 µM and 30 µM being significantly different from vehicle (n = 7, one-way ANOVA, p > 0.001, Figure 1B and 1D, open-circles). CP55940 hyperpolarised AtT20-CB_1_ cells with a *p*EC_50_ of 7.64 ± 0.08 (n_H_ 1.2 ± 0.3) to a maximum of 34.3 ± 1.6 % (Figure 1B and Figure 1D, closed-diamonds). (+)-CBD (1-30 µM) did not produce a change in fluorescence in AtT20-WT cells that was different from vehicle-controls (n = 11, one-way ANOVA p = 0.32, Figure 1C and 1D, grey closed-circles).

The effects of (+)-CBD (10 µM) were significantly inhibited by pretreatment of AtT20-CB_1_ or AtT20-CB_2_ cells with pertussis toxin (200 ng.mL^−1^ overnight, p < 0.05 for each, n = 7; Figure 2).

**Figure 2.**
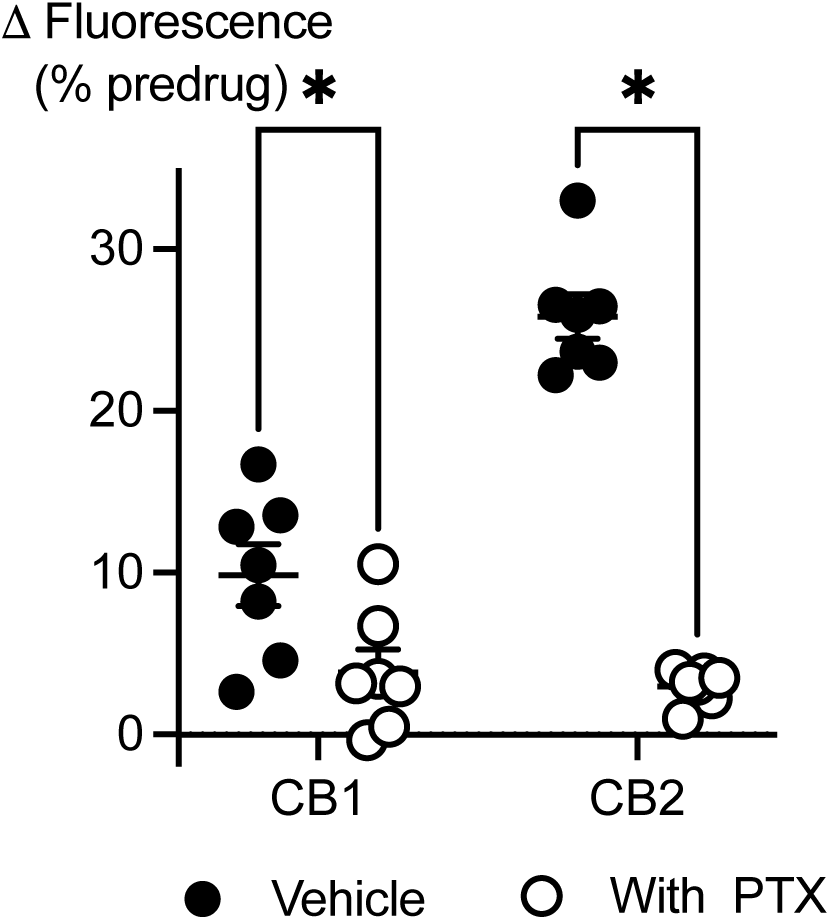
The effects of (+)-CBD are mediated via pertussis toxin-sensitive G proteins. AtT20 cells expressing CB_1_ or CB_2_ receptors were challenged with (+)-CBD (30 µM) after overnight pretreatment with pertussis toxin (PTX, 200 ng mL^−1^) or vehicle. The effects of (+)-CBD were significantly (*) reduced by PTX (Student’s unpaired t-test, p < 0.05). Each dot represents one independent experiment performed in duplicate; bars indicate mean ± SEM (n = 7).

### Quantification of CB_2_ Receptor Antagonism

To further explore the interaction of (+)-CBD at CB_2_ receptors, we conducted a Schild analysis using the CB_2_ receptor antagonist AM630 and compared the effects of AM630 on (+)-CBD and its effects on the orthosteric agonist CP55940. Increasing concentrations of AM630 produced a similar parallel shift in the concentration response curves to both (+)-CBD (Schild slope 1.1 ± 0.1, (Figure 3) and CP55940 (Schild slope 0.9 ± 0.1), the *p*A_2_ for AM630 was 6.5 ± 0.16 for (+)-CBD and 6.0 ± 0.17 for CP55940.

**Figure 3.**
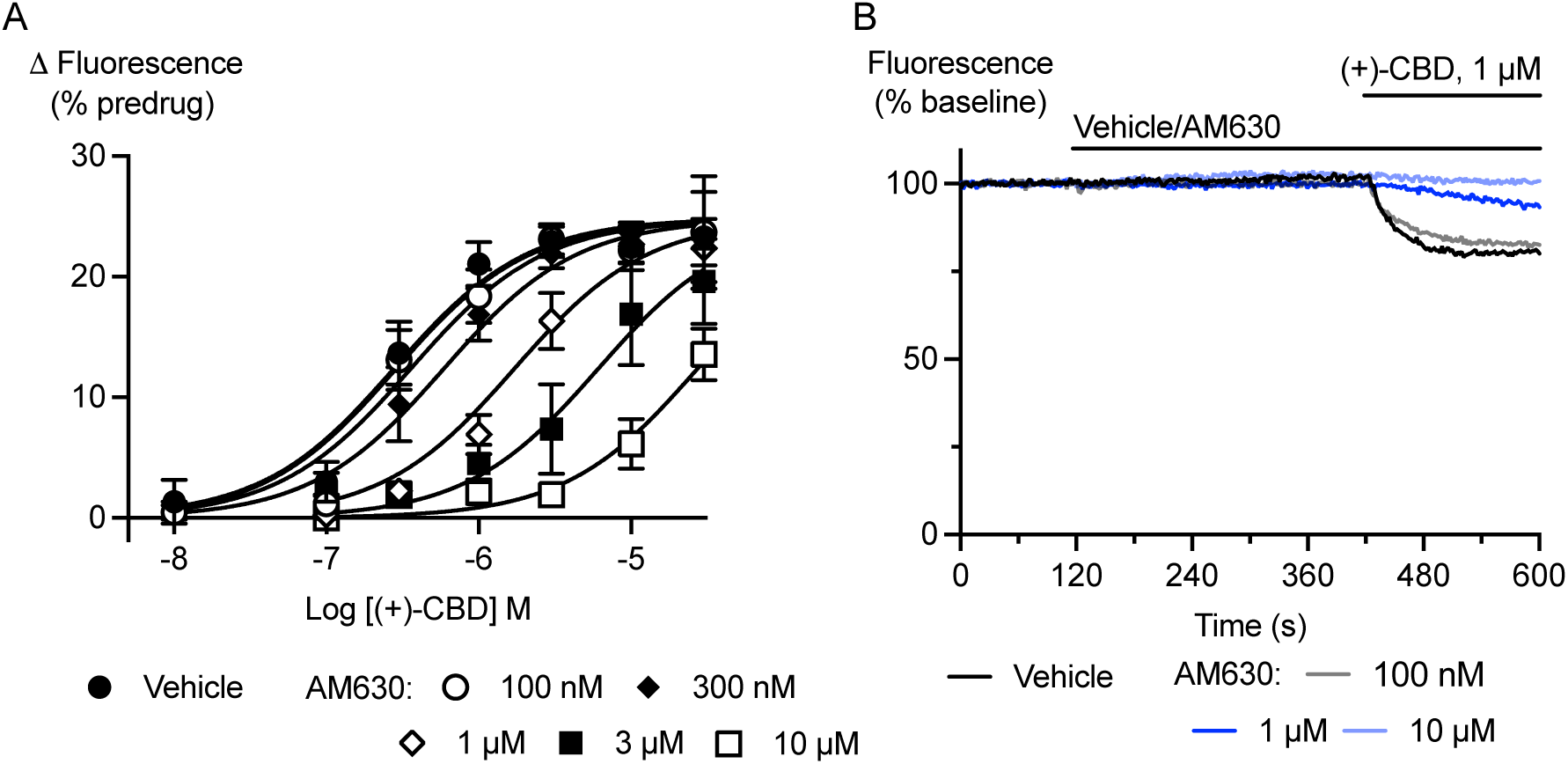
The effects of (+)-CBD are inhibited by the CB_2_ antagonist AM630. AtT20 cells expressing CB_2_ receptors were preincubated with AM630 (100 nM to 10 µM) and then challenged with (+)-CBD (10 nM – 30 µM). **A)** Represents the pooled concentration response curves for (+)-CBD in the presence of the indicated concentrations of (+)-CBD. The curves were shifted in parallel as AM630 concentrations increased (Schild slope 1.1 ± 0.1, *p*A_2_ 6.5 ± 0.2, n = 6 independent experiments). **B)** Example traces showing the progressive reduction in the response to (+)-CBD (1 µM) produced by increasing concentrations of AM630.

### CB_1_ Receptor Antagonism

We examined whether a selective CB_1_ antagonist, SR141716A, blocks (+)-CBD signal on human CB_1_. A 5-minute pretreatment of AtT20-CB_1_ cells with SR141716A (3 µM) significantly reduced the signal of 10 µM (+)-CBD from 6.5 ± 1.1 % to 1.1 ± 0.7 % (p < 0.05, n = 7, Figure 4A). This pretreatment also blocked the effects of 100 nM CP55940, a submaximal effective concentration, from 25.5 ± 1.4% to -2.0 ± 1.1 % (p < 0.05, n = 7, Figure 4A).

**Figure 4.**
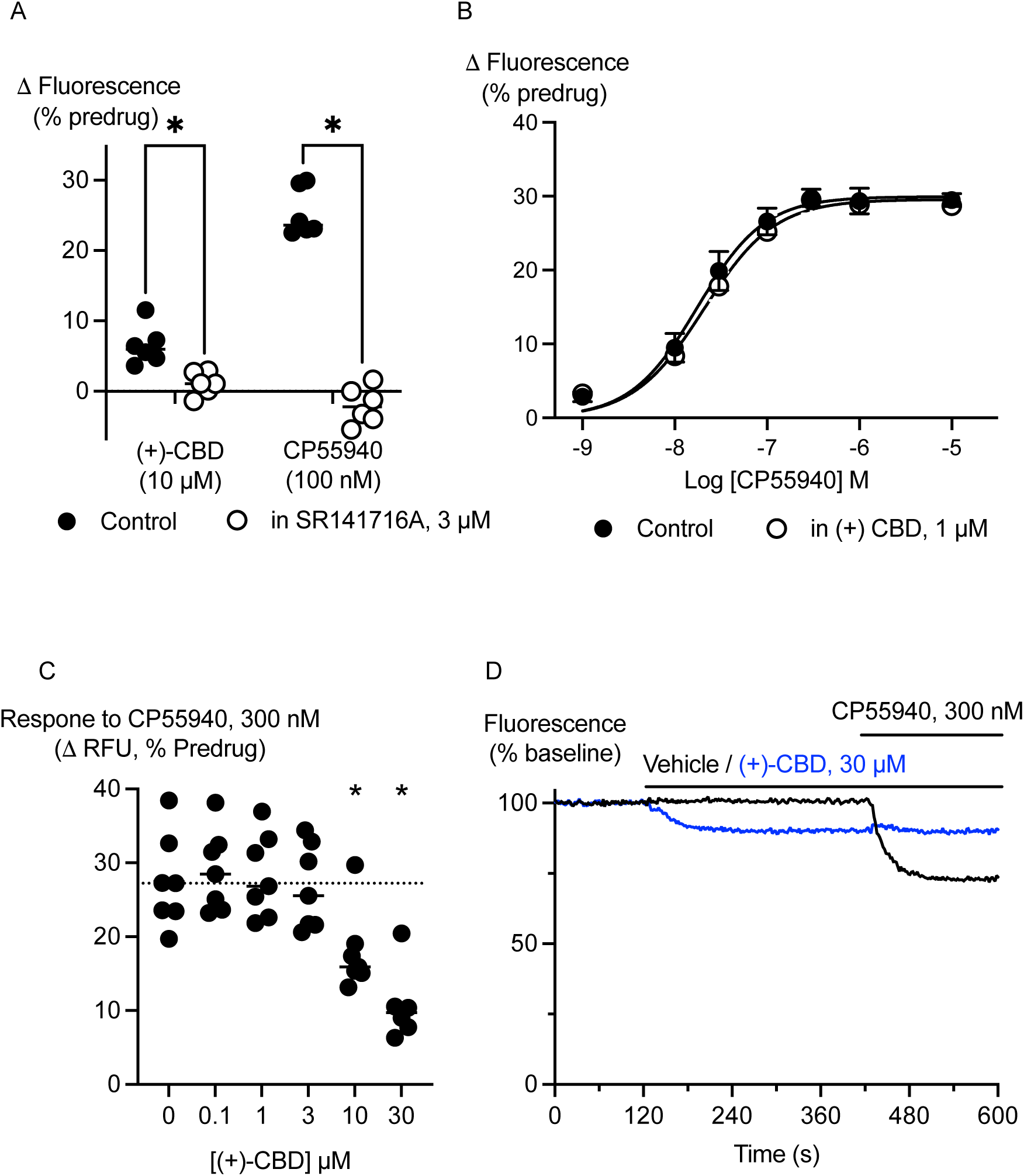
(+)-CBD is a low efficacy, low potency agonist of CB_1_ receptors. AtT20 cells expressing CB_1_ receptors were preincubated for 5 minutes with (+)-CBD or SR141716A and challenged with (+)-CBD or CP55940. **A)** Preincubation with SR141716A (3 µM) significantly (*) inhibited the effects of (+)-CBD (30 µM) and CP55940 (100 nM; Student’s unpaired t-test, p < 0.05 for each). **B)** Preincubation with (+)-CBD, 1 µM, did not affect the concentration response relationship to subsequent application of CP55940 (n = 6). **C)** preincubation with (+)-CBD (10 µM, 30 µM) significantly (*) inhibited the effects of CP55940 (300 nM; one-way ANOVA, p < 0.0001, followed by Dunnett’s test for multiple comparisons, n = 7). **D)** Example traces showing the hyperpolarisation of AtT20-CB_1_ cells by (+)-CBD, and the subsequent inhibition of the response to CP55940.

(+)-CBD has previously been identified as a CB_1_ antagonist in mouse neurons (21). We determined that high concentrations of (+)-CBD (10 µM, 30 µM) produced a significant inhibition of responses to a high concentration of CP55940 (300 nM) (n = 7, one-way ANOVA p > 0.001, Figure 4C and 4D). However, pretreatment of AtT20-CB_1_ cells with a concentration of (+)-CBD (1 µM) that produced no change in membrane potential by itself had no effect on the response to subsequently applied CP55940 as shown in Figure 4B (CP55940 alone; *p*EC_50_ 7.75 ± 0.08, maximum response 29.9 ± 1.1, n_H_ 1.2 ± 0.2; CP55940 in the presence of 1 µM (+)-CBD; 7.68 ± 0.08, maximum response 29.6 ± 1.2, n_H_ 1.1 ± 0.2, n = 6)

### Other effects of (+)-CBD

In AtT20-CB_1_ cells, the apparent effects of (+)-CBD on responses to a high efficacy agonist could arise from actions at the CB_1_ receptor or actions at the signalling effector, G protein-gated inwardly rectifying K channels (GIRK channels). We tested the latter possibility in two ways: by pre-incubating AtT20-WT cells with (+)-CBD (1-30 µM) and then activating GIRK channels via endogenous somatostatin receptors (SSTR), and by directly activating GIRK channels with ML297 following pre-incubation with (+)-CBD. (+)-CBD (10 µM, 30 µM) inhibited the hyperpolarisation produced by 100nM somatostatin-14 (SRIF-14; n = 6, one-way ANOVA p = 0.0027, Figure 5A and 5B) and by 30 µM ML297 (one-way ANOVA p = 0.025, Figure 5C and 5D). At 30 µM, (+)-CBD inhibited the activation of GIRK by SRIF-14 by 41 ± 4 % and the direct activation of GIRK by ML297 by 14 ± 2 %, suggesting that (+)-CBD directly inhibits SSTR in addition to GIRK.

**Figure 5.**
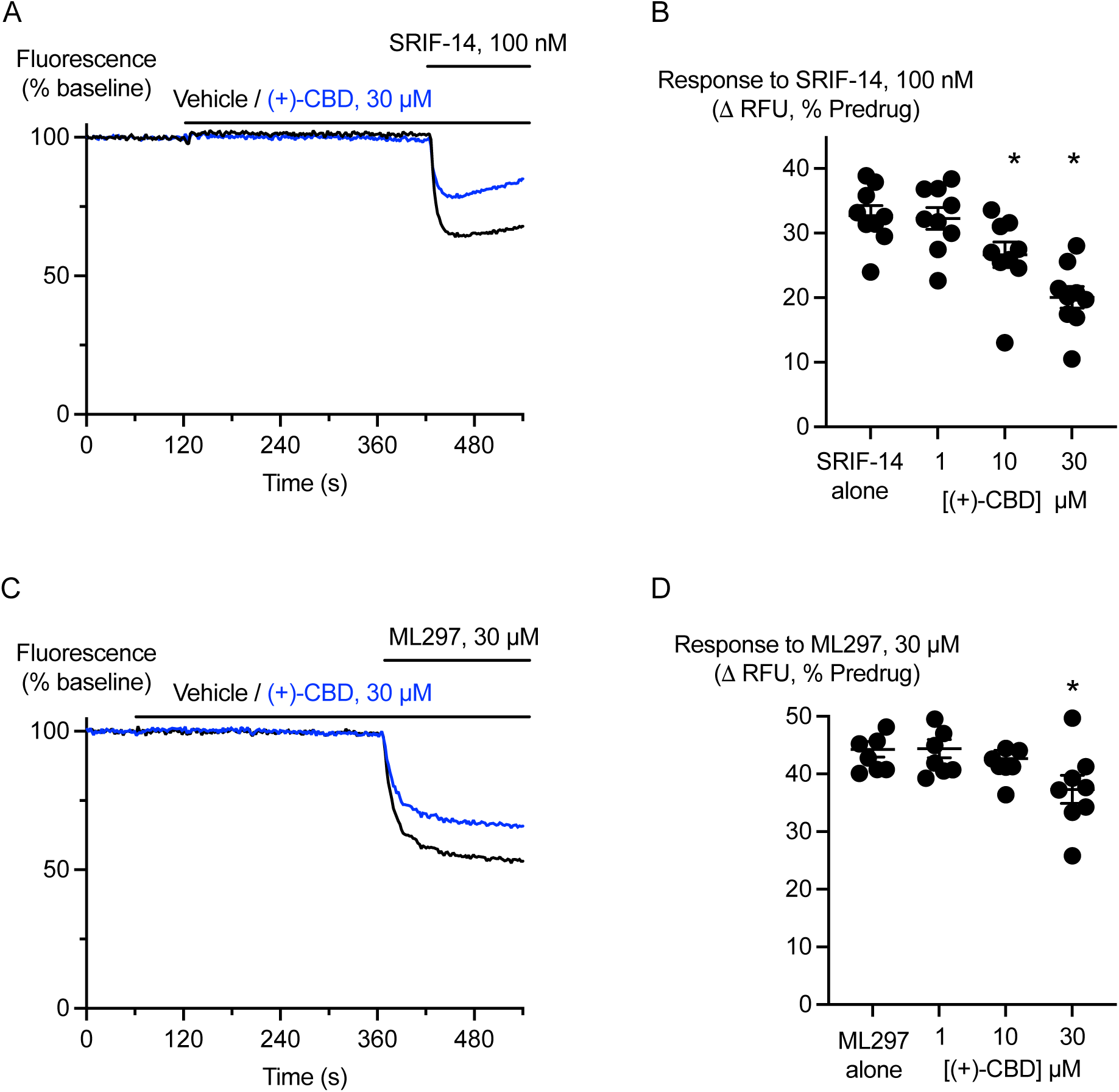
(+)-CBD is a low potency inhibitor of somatostatin receptors and GIRK channels. AtT20-WT cells (expressing neither CB_1_ or CB_2_ receptors) were preincubated with (+)-CBD for 5 minutes and challenged with SRIF-14 (100 nM) or the GIRK activator ML297 (30 µM). **A)** Example traces showing that (+)-CBD (30 µM) did not change the membrane potential of AtT20-EV cells but inhibited their hyperpolarisation by SRIF-14. **B)** A summary plot of the effects of (+)-CBD on the responses to SRIF-14 (100 nM), (+)-CBD (10 µM, 30 µM) significantly (*) inhibited the responses (one-way ANOVA, p < 0.0001, followed by Dunnett’s test for multiple comparisons, n = 8). **C)** Example traces showing that (+)-CBD (30 µM) inhibited the hyperpolarisation of AtT20-WT cells by ML297. **D)** A summary plot of the effects of (+)-CBD on the responses to ML297 (30 µM), (+)-CBD (30 µM) significantly (*) inhibited the responses (one-way ANOVA, p = 0.0253, followed by Dunnett’s test for multiple comparisons, n = 8).

In contrast to (+)-CBD, 5 minute application of (−)-CBD (10 µM, 30 µM) produced a small apparent hyperpolarisation of AtT20-WT cells (n = 11, one-way ANOVA p < 0.0001, Supplementary Figure 2A) but did not inhibit the hyperpolarisation of AtT20-WT cells by 100 nM SRIF-14 (n = 8, one-way ANOVA p = 0.8344, Supplementary Figure 2B). (−)-CBD (30 µM) also failed to inhibit the hyperpolarisation produced by 30 µM ML297 (n = 7, one-way ANOVA p = 0.3869, Supplementary Figure 2C).

### (+)-CBD and (−)-CBD docking in human CB_2_ receptor

Molecular docking simulations were conducted to investigate the differences in how the enantiomers of CBD interact with the CB_2_ receptor (Fig. 6). The results revealed a notable difference in docking scores: (+)-CBD exhibited a more favourable binding affinity (-8.224 kcal/mol) compared to (−)-CBD (–7.414 kcal/mol). A key distinction in their binding modes was observed in the interaction with the amino acid residue Ser285. In the top-ranked binding pose, one of the hydroxyl groups of (+)-CBD formed a moderately strong hydrogen bond with Ser285, with an oxygen–oxygen (O–O) distance of 2.88 Å. In contrast, (−)-CBD did not closely approach this residue, maintaining an O–O distance of 3.70 Å, which suggests little to no hydrogen bonding with Ser285. For comparison, the synthetic cannabinoid CP55940 showed an even more favourable docking score (–8.99 kcal/mol) and was found to engage more extensively with lipophilic residues within the receptor pocket. Notably, it formed a strong hydrogen bond with Ser285, with an O–O distance of 2.59 Å.

**Figure 6.**
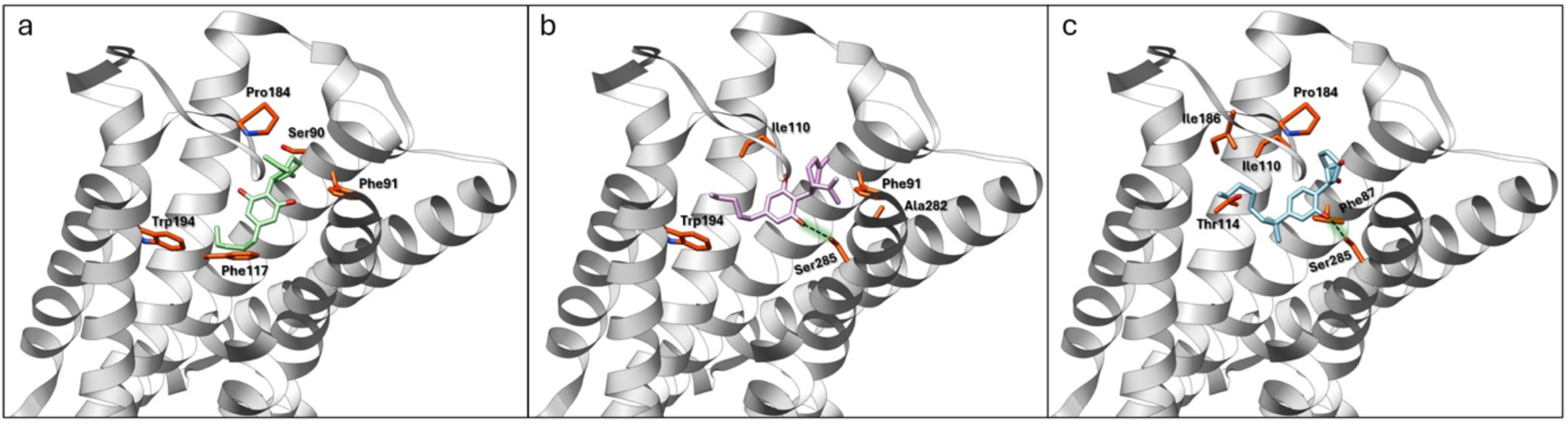
Top energetically favourable 3D binding poses of (−)-CBD (a), (+)-CBD (b), and CP55940 (c) inside the CB2 receptor pocket. Some key surrounding amino acids stabilizing the interaction of the CB2 receptor with the ligand are highlighted in orange. H-bonds between Ser285 and ligands are shown by dashed black lines in b and c. Some residues are omitted for clarity.

## Discussion

The principal finding of this study is that (+)-CBD is a robust agonist at human CB_2_ receptors, in sharp contrast to (−)-CBD, which has little CB_2_ activity under these conditions. The effects of (+)-CBD were inhibited by the CB_2_ antagonist AM630, blocked by pretreatment of cells with pertussis toxin and (+)-CBD had no effect on the membrane potential of AtT20-FlpIn cells that were not engineered to express CB_2_. (+)-CBD has been reported to have an affinity of 200 nM for CB_2_ (species unspecified) in a radioligand binding assay (19,41), which is similar to the EC_50_ we observed in the assay of cellular membrane potential (235 nM). We are unaware of any other reports of (+)-CBD activity at CB_2_ receptors in vitro. One limitation of the assay we used is that (+)-CBD modestly (14%) inhibited GIRK channel activation by ML297 at the highest concentrations tested, this may explain the reduced maximum of CBD compared to the high efficacy agonist CP 55940 (about 10% less). We were also unable to obtain the other 2 isomers of CBD – cis**-**(3*R*,4*S*)**-**CBD and cis-(3S,4R)-CBD for testing.

In vivo, (+)-CBD and analogues have been reported to be anticonvulsant and inhibit ovulation in rodent models; however, the receptor(s) involved in these effects were not identified (42,43). CBD has been reported to have weak agonist activity in an assay of cAMP accumulation in CHO cells expressing CB_2_ (EC_50_ > 10 µM)(12), potential allosteric antagonist activity in an assay of GTPγS binding in CB_2_ receptor expressing CHO cells (10) and in cAMP accumulation assays in HEK293 cells expressing CB_2_ (13). All these experiments appear to have been carried out with plant-derived, presumably (−)-CBD. A study by Moniruzzaman and colleagues (44) reported that synthetic CBD inhibited the proliferation and migration of several colon cancer cell lines in a manner was sensitive to the CB_2_ antagonist SR144528, and in one case the CB_1_ antagonist AM251. The concentrations of CBD used was quite high (5 µg.mL^−1^ minimum, or about 16 µM), but the results are consistent with what is reported here if the synthetic CBD was a racemic mixture or even pure (+)-CBD.

(+)-CBD had modest activity at CB_1_ and after brief pre-incubation it diminished the effects of CP55940, consistent with a moderate affinity and low efficacy at this receptor. At rat CB_1_, (+)-CBD has been reported to have an affinity of 250 nM-840 nM (19,21,41) and it acts as a potent antagonist of mouse CB_1_ activation in assays of autaptic neurotransmission (21).

Under the same conditions in which (+)-CBD was an effective CB_2_ agonist, (−)-CBD had no acute agonist activity and little effect on CP55940 signalling at either CB_1_ or CB_2_; consistent with its activity as a negative allosteric modulator of CB_1_ (10,11). The effects of (−)-CBD in cell signalling assays are complex, which may in part reflect specific interactions with a wide-variety of G protein-coupled receptors (and other proteins) as well as unspecific actions caused by effects on cell membranes or even colloidal aggregation of (−)-CBD (20,45). We are confident that the agonist activity of (+)-CBD at CB_2_ was specific for a number of reasons. The effect was sensitive to the generally accepted inhibitor of orthosteric CB_2_ receptor function, the CB_2_ antagonist AM630. Receptor signalling was significantly reduced by pertussis toxin (PTX), an inhibitor of G_i_/G_o_-type G proteins which are the primary transducers of CB_2_ signalling; and (+)-CBD had no effect on membrane potential in AtT20 cells not expressing cannabinoid receptors. Noteworthy, the concentration of (+)-CBD that activated CB_2_ receptors was well below those reported to result in aggregation of (−)-CBD (45,46) and consistent with the reported affinity of (+)-CBD for CB_2_ (Bisogno). The n_H_ for activation of CB_2_ by (+)-CBD was similar to that of CP55940, and not different from 1 – unspecific effects on receptor activation produced by drugs which aggregate in solution often display steep nH (45).

(+)-CBD appeared to be a low efficacy, low potency agonist at CB_1_. The hyperpolarisation produced by (+)-CBD was sensitive to the CB_1_ antagonist SR141716A and pertussis toxin. High concentrations of (+)-CBD reduced the activation of CB_1_ by CP55940, consistent with it being a low efficacy agonist. High concentrations of (+)-CBD (but not (−)-CBD) also inhibited somatostatin-induced hyperpolarisation of AtT20 cells, suggesting weak antagonist activity at the endogenous SSTR in AtT20 cells.

Molecular docking simulations suggest that the difference in CB_2_ receptor activity between (−)-CBD and (+)-CBD is primarily driven by differences in binding affinity and enhanced interaction with a residue important for CB_2_ activation. (+)-CBD, which exhibited a more negative docking score, is predicted to form more energetically favourable interactions with CB_2_ and thus demonstrates higher affinity. Similarly, CP55940 showed an even more favourable docking score than both CBD enantiomers, consistent with its experimentally observed high binding affinity for CB_2_ (30). A key factor contributing to the differential affinity appears to be the ability of the compounds to form hydrogen bonds with the serine residue at position 285 (Ser285). Previous mutagenesis studies have shown that Ser285 plays a critical role in ligand binding and agonist activity for some cannabinoid-like compounds, including CP55940 (30). Both CP55940 and (+)-CBD form hydrogen bonds with Ser285, with CP55940 exhibiting a particularly strong interaction, indicated by a short O–O distance of 2.59 Å (Fig 6c). (+)-CBD also forms a moderately strong hydrogen bond with Ser285 (O–O distance of 2.88 Å)(Fig 6b). In contrast, (−)-CBD, due to its enantiomeric configuration, fails to position its phenolic oxygen close enough to Ser285 to form a meaningful interaction (Fig 6a). With an O–O distance of 3.70 Å, simulations suggest only a weak or negligible hydrogen bond between (−)-CBD and Ser285. These findings indicate that the enantiomeric configuration in the CBD structure can significantly influence how different functional groups orient within the CB_2_ receptor binding pocket and directly impact receptor affinity

To our knowledge only one study has reported the effects of administration of (+)-CBD to humans (23). That study used robust doses of (+)-CBD (200 mg – 800 mg) administered before a cold pressor test to measure effects on pain thresholds and pain tolerance when compared with placebo. There were small or insignificant effects of (+)-CBD on acute pain threshold or tolerance, although all doses increased the perceived painfulness of the stimulus. (+)-CBD did not affect the subjective mood or the heart rate of participants, consistent with the limited the activity of (+)-CBD at CB_1_ we measured, but it did produce a small but significant decrease in blood pressure. Overall, (+)-CBD was well tolerated, similar to other CB_2_ agonists trialled in humans. The limited effect of acute administration of a CB_2_ agonist on pain responses is perhaps not surprising, given the limited expression of CB2 receptors in human sensory neurons, and the lack of evidence of expression on central neurons involved in pain responses (in humans). It should be noted that many clinical trials using CBD do not specify the molecular identity of the drug, and it is possible that trials using synthetic CBD are administering either racemic mixtures of CBD, or indeed simply (+)-CBD.

This study emphasises the importance of understanding the molecular actions of the enantiomers of chiral cannabinoids. Unlike THC and CBC, there does not seem to be any evidence for the occurrence of (+)-CBD in *Cannabis* plants, but synthetic CBD may have reasonably robust CB_2_ agonist activity if it contains a substantial amount of (+)-CBD. Although the safety of (+)-CBD cannot simply be assumed to be the same as (−)-CBD, the identification of (+)-CBD as a robust CB2 agonist suggests that (+)-CBD may be worth exploring as a therapeutic agent in chronic inflammatory conditions or other diseases where CB_2_ activation may prove beneficial.

## Acknowledgements

We thank A/Prof Alf Garcia-Barnett and Prof Alison Rodger for help with the CD determinations. This work was supported by recurrent research funding to MC and by the Australian Research Council Industrial Transformation Center for Facilitated Advancement of Australia’s Bioactives (Grant IC210100040) and Research Attraction and Acceleration Program funding from the Office of the Chief Scientist and Engineer, Investment NSW.

## Data Availability

Original data is available upon reasonable request.

## Author Contribution Statement

**PBB:** Investigation (lead); formal analysis; writing - original draft (supporting), writing - review and editing. **CC**: investigation; formal analysis, writing - review and editing. **MH**: Investigation; formal analysis; writing - original draft (supporting); writing – review and editing. **EJMC:** Investigation; formal analysis. **DM**: Investigation; formal analysis. **MJS**: Supervision (equal); conceptualisation, investigation; methodology; formal analysis; writing – original draft (supporting); writing – review and editing (equal). **MC** Conceptualisation (lead); investigation, formal analysis, visualisation; writing – original draft (lead); supervision (equal)

## Abbreviations

CBC: cannabichromene
CD: circular dichroism
(−)-CBD: (−)-*trans*-cannabidiol
(+)-CBD: (+)-*trans*-cannabidiol
GIRK: G protein-gated inwardly rectifying K channels
HBSS: modified Hank’s buffered saline solution
MPA: membrane potential assay
PTX: pertussis toxin
SSTR: somatostatin receptors
SRIF-14: somatostatin-14
THC: Δ^9^-tetrahydrocannabinol

---

**Supplementary Figure 1.**
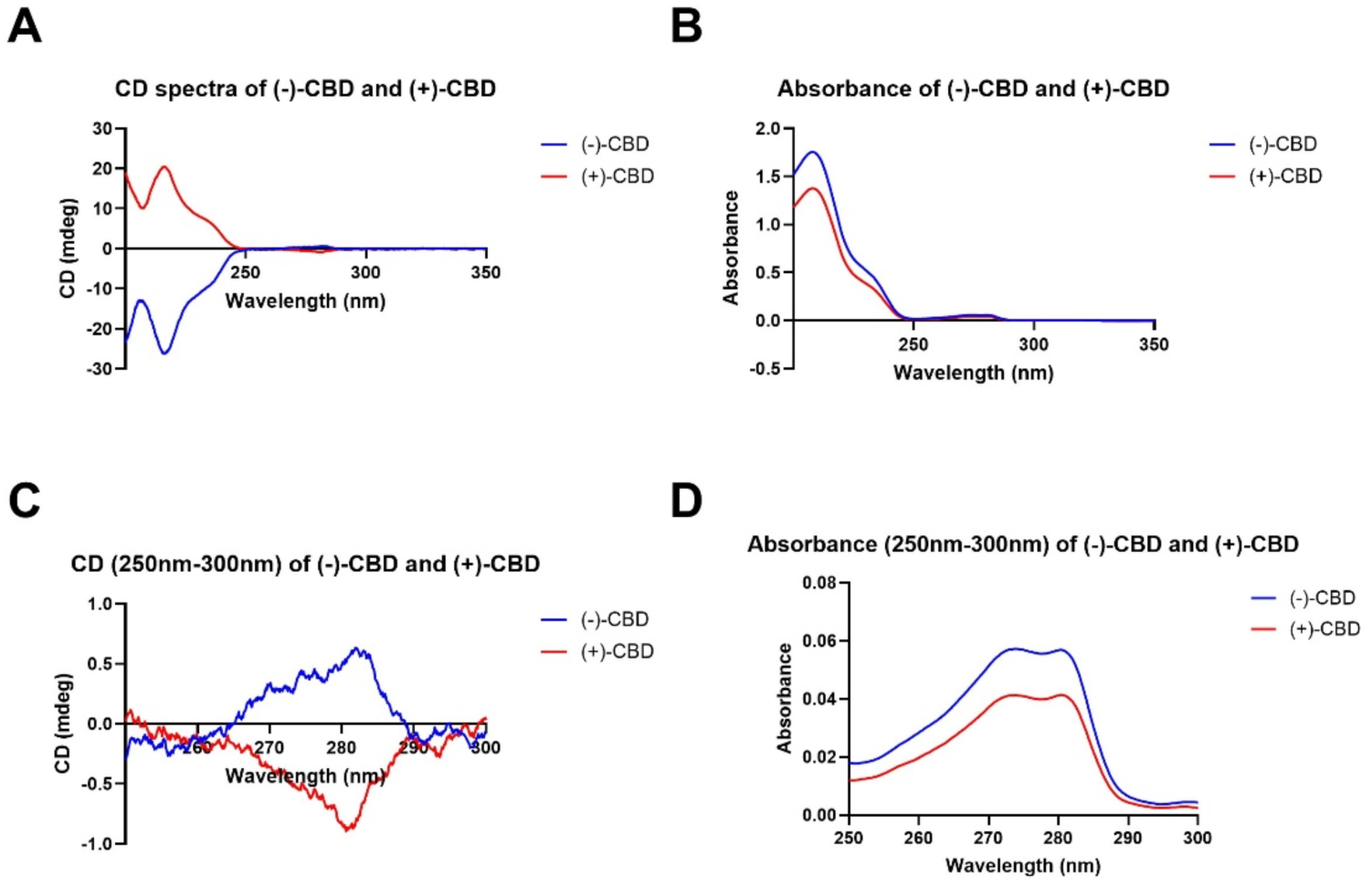
(−)-CBD and (+)-CBD used in this study have distinct optical spectra. The absorbance and circular dichroism (CD) spectra of the samples were determined on a Jasco (Hachioji, Japan) J-1500 spectropolarimeter by diluting the enantiomerically pure sample to 0.1 mg mL^−1^ in ethanol, with 0.1 mm path length. (A and C) CD spectra and (B and D) absorbance spectra of (−)-CBD and (+)-CBD.

**Supplementary Figure 2.**
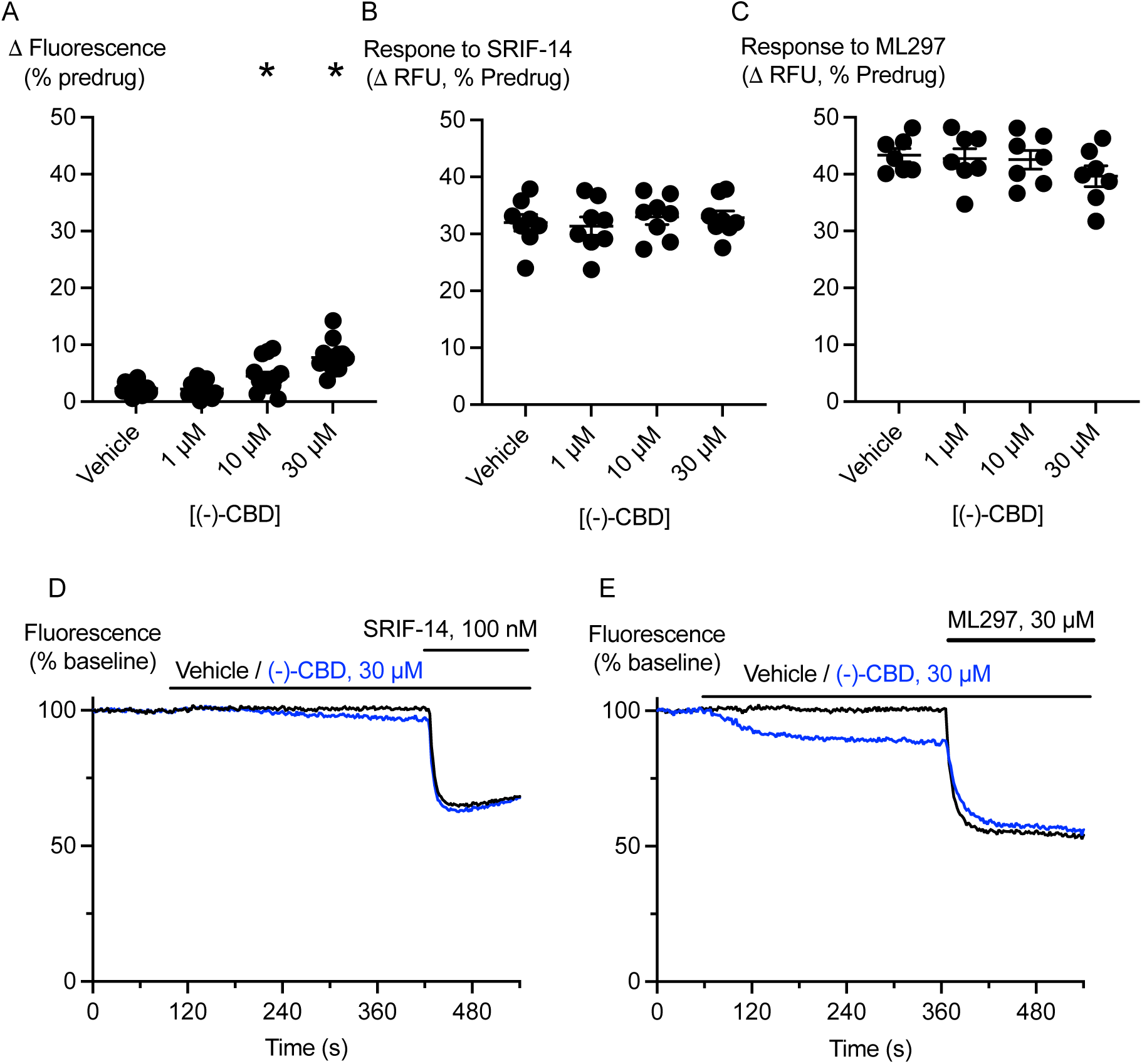
(−)-CBD has distinct effects to (+)-CBD in cells not expressing cannabinoid receptors. AtT20-WT cells were incubated with (−)-CBD for 5 minutes and then challenged with SRIF-14 (100 nM) or the GIRK activator ML297 (30 µM). **A)** A summary plot of the effects of (−)-CBD alone on the fluorescence of AtT20-WT cells, (−)-CBD (10 µM, 30 µM) produced a significant (*) apparent hyperpolarisation of the cells compared to vehicle (one-way ANOVA, p = 0.0001, followed by Dunnett’s test for multiple comparisons, n = 15). **B)** (−)-CBD did not inhibit the response of AtT20-WT cells to SRIF-14 (100 nM) (one-way ANOVA, p = 0.8344, n = 8). **C)** (−)-CBD did not inhibit the response of AtT20-WT cells to ML297 (30 µM) (one-way ANOVA, p = 0.3869, n = 7). **D)** Example traces showing the effect (−)-CBD (30 µM) on SRIF-14 responses. **E)** Example traces showing the effect (−)-CBD (30 µM) on ML297 responses.

Reagents

(−)-CBD, NMI, # D512c

(+)-CBD, Cayman Chemical, # 9003416

(−)-CP55,940, Cayman Chemical, # 90084

AM630, Cayman Chemical, # 10006974

SR141716A, Cayman Chemical, # 9000484

Somatostatin1-14, AUSPEP, # 2076

Pertussis Toxin, Hello Bio #HB4729

